# Efficient Inhibition of SARS-CoV-2 Using Chimeric Antisense Oligonucleotides through RNase L Activation

**DOI:** 10.1101/2021.03.04.433849

**Authors:** Xiaoxuan Su, Wenxiao Ma, Boyang Cheng, Qian Wang, Zefeng Guo, Demin Zhou, Xinjing Tang

**Author notes:** Corresponding author: Xinjing Tang. These authors contributed equally to this work (Xiaoxuan Su, Wenxiao Ma).

## Abstract

There is an urgent need for effective antiviral drugs to alleviate the current COVID-19 pandemic. Here, we rationally designed and developed chimeric antisense oligonucleotides to degrade envelope and spike RNAs of SARS-CoV-2. Each oligonucleotide comprises a 3’ antisense sequence for target recognition and a 5’-phosphorylated 2’-5’ poly(A)4 for guided ribonuclease L (RNase L) activation. Since RNase L can potently cleave single strand RNA during innate antiviral response, the improved degradation efficiency of chimeric oligonucleotides was twice as much as classic antisense oligonucleotides in Vero cells, for both SARS-CoV-2 RNA targets. In pseudovirus infection models, one of chimeric oligonucleotides targeting spike RNA achieved potent and broad-spectrum inhibition of both SARS-CoV-2 and its recently reported N501Y and/or ΔH69/ΔV70 mutants. These results showed that the constructed chimeric oligonucleotides could efficiently degrade pathogenic RNA of SARS-CoV-2 facilitated by immune activation, showing promising potentials as antiviral nucleic acid drugs for COVID-19.

## Introduction

Since the infection was first reported in 2019, severe acute respiratory syndrome coronavirus 2 (SARS-CoV-2) has continued to spread globally and caused the pandemic COVID-19 disease (*1*). The current lack of highly effective antiviral drugs for SARS-CoV-2 has made the treatment of infected patients more difficult, thus demanding more candidate options for drug discovery. Genomic positive-sense single-stranded RNA (ssRNA) and structural proteins participate in virus packaging, which is an essential step in SARS-CoV-2 life cycle. Envelope (E), spike (S) and membrane (M) proteins assemble the virus membrane in host cells infected by SARS-CoV-2 (*2, 3*), and thus become ideal drug targets to intervene virus proliferation.

RNase L participates in innate antiviral response of vertebrate cells by cleaving UN^N sites located in viral or cellular ssRNAs. Cytoplasmic RNase L monomer only displays weak catalytic cleavage on the substrate. However, upon dimerization induced by its specific ligand 5’ phosphorylated 2’5’ polyA (such as 4A_2-5_), RNase L is highly activated and performs intense RNA cleavage (*4*). The cleavage products can further bind to intracellular pattern recognition receptors (PRRs) to stimulate the production of interferons (IFN) (*5–7*), which in turn induces the expression of interferon stimulated genes (ISGs) including RNase L, to enhance the antiviral response (*2, 7, 8*). Ubiquitous activation of RNase L might cause widespread attenuation of basal mRNA and possible cell apoptosis, especially at high doses of 4A_2-5_ (*9–12*). Guided and controlled activation of RNase L could otherwise achieve more specific target RNA degradation. RNA binding small molecules conjugated with 4A_2-5_ have been reported to target highly-structured microRNA or RNA fragments of virus genome (*13, 14*), which contains particularly structured sequences. Nevertheless, the selective binding between a small molecule and the specific region of pathogenic RNA is limited, while the sequence-selective antisense oligonucleotides (ASO) will be more accessible and effective to target viral RNA of interest.

ASO therapy has successfully targeted undruggable pathogenic genes of rare diseases and has been developed against the infection of ssRNA viruses such as SARS-CoV (*15*) in a sequence-specific manner. Chemical modifications on ASOs can further promote their nuclease resistance and/or binding affinity to target RNA sequences, such as phosphorothioate (PS) linkages and 2’-*O*-methyl (2’-OMe) substituents (*16*). Currently a few reports have raised the possibility of combining ASOs with 4A_2-5_ for the treatment of tumors (*17*) and viral infections (*12, 18*). Therefore, it is promising to develop nucleic acid drugs in form of ASO-4A_2-5_ chimera targeting SARS-CoV-2 genomic RNAs to inhibit virus infection.

Here, based on nucleic acid-hydrolysis targeting chimeras strategy, we developed chimeric antisense oligonucleotides with 4A_2-5_ conjugation through flexible PEG linker to target envelope RNA (Chimera-E) or spike RNA (Chimera-S) of SARS-CoV-2. The antisense component specifically recognizes complementary target RNA sequence, while the covalently linked 4A_2-5_ moiety functions as RNase L recruiter, thus collectively guiding RNase L to specific cleavage sites on targeted viral RNA. With these ASO-4A_2-5_ chimeras, we evaluated RNA knockdown of both SARS-CoV-2 envelope and spike genes in Vero cells. Further in a pseudotyped SARS-CoV-2 infection model, ASO-4A_2-5_ chimeras for spike gene successfully inhibited pseudovirus packaging and further infection on host cells. One of these chimeras targeting spike gene also effectively inhibited three mutants of SARS-CoV-2 pseudovirus, including N501Y, ΔH69/ΔV70, and the recently discovered dual-site mutations with higher spreading ability. In addition, these chimeras could upregulate the expression of RNase L and cytokines (such as IFN-β and IL-6) as antiviral immune responses *in vitro.* The antiviral efficacy and versatility of the 4A_2-5_-modified chimeric oligonucleotides provide a new treatment option for the current COVID-19 pandemic.

## Results

### Rational design and characterization of RNase L-recruiting chimeric antisense oligonucleotides

Our study began with the selection of antisense oligonucleotides targeting specific genomic RNA of SARS-CoV-2. After predicting RNA secondary structures of spike receptor binding domain (S-RBD) and envelope (E) protein of SARS-CoV-2, loops composed of more than 10 nucleotides were selected as ideal target regions. In addition, considering the space required for RNase L activation and substrate cleavage, the stem structure in 3’ proximity of the selected loop was limited to have less than 4 base pairs, and its 3’ pairing end should have more than 1 RNase L cleavage site (UN^N) in a bulge structure. As a result, antisense sequences complementary to the selected loops were predicted with more than 70% probability of being efficient antisense strands as evaluated by OligoWalk (*19*) and was synthesized through solid phase synthesis (**Table S1**).To enhance nuclease resistance and binding affinity with their complementary viral RNA regions, phosphorothioate (PS) linkages and 2’-*O*-methyl (2’-OMe) substituents were properly incorporated into the chimeric structure, followed by the coupling of a poly 2’-5’ poly(A)4 ligand at 5’ terminus of the designed antisense sequence (15 nt) through a short PEG linker (**Fig. 1B**).

**Fig 1.**
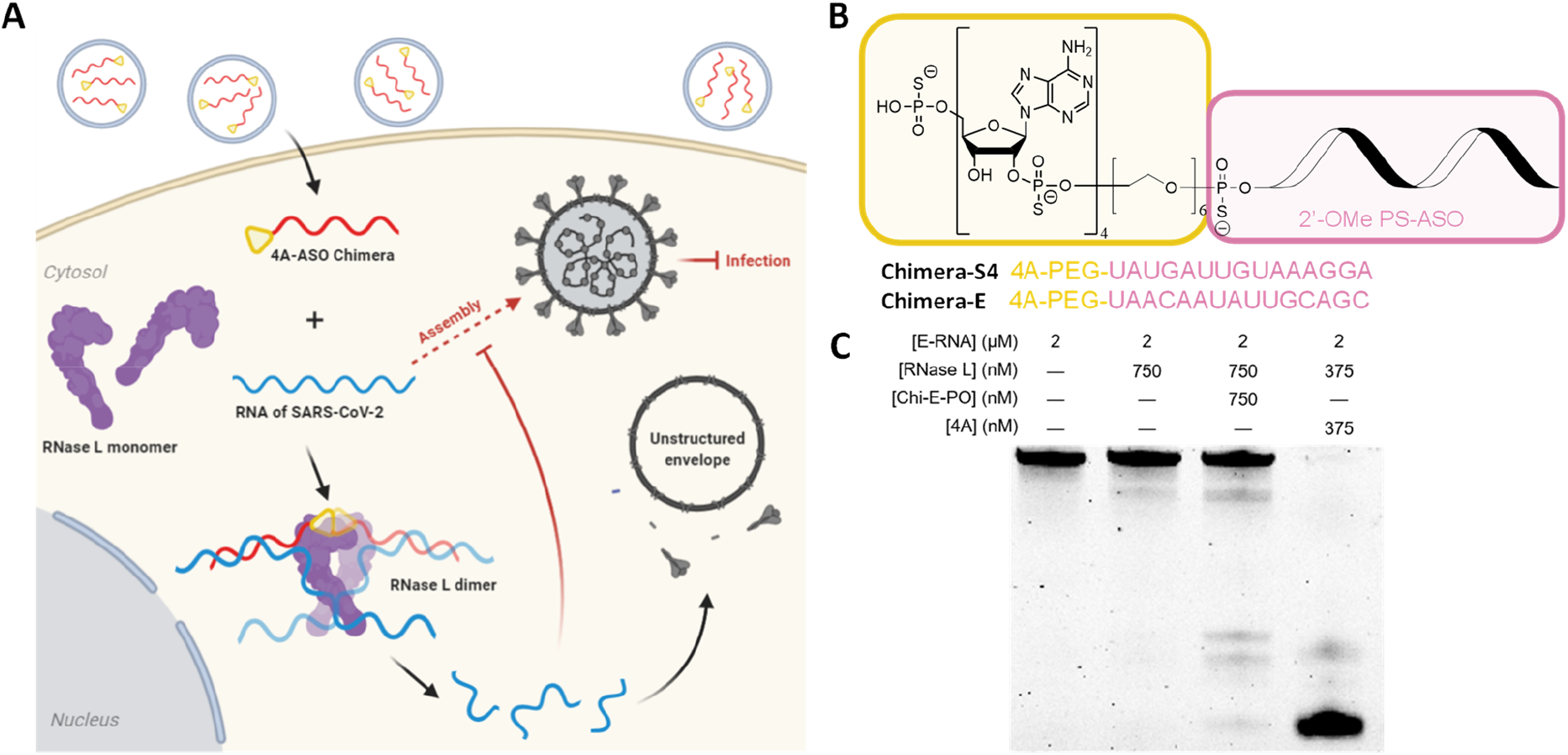
Rational design of 4A_2-5_-ASO chimeric antisense oligonucleotides to target and degrade SARS-CoV-2 RNA by RNase L recruitment. **(A)** Schematic representation of 4A_2-5_-ASO chimera induced inhibition of SARS-CoV-2 proliferation. Black arrows, the actual processing of viral RNA upon treatment; dashed arrow, inhibited viral assembly during SARS-CoV-2 infection. **(B)** Structures of 4A_2-5_-ASO Chimeras targeting envelope-(E-) and spike- (S-) RNA of SARS-CoV-2, respectively. **(C)** *In vitro* cleavage assay of a 3’ Cy3-labeled E-RNA segment (62 nt).

We first tested RNase L recruitment ability of Chimera-E-PO, an oligonucleotide modified with 5’ native 4A_2-5_ ligand and complementary to a loop structure on Cy3-labeled partial E-RNA sequence of SARS-CoV-2 (**Table S1**). As shown in **Fig 1C**, after incubating RNase L with Chimera-E-PO or 4A_2-5_, *in vitro* cleavage of Cy3-labeled substrate RNA was analyzed in a denaturing PAGE gel. Treatment of RNase L alone did not lead to the cleavage of substrate RNA, while additional Chimera-E-PO treatment activated RNase L and produced cleavage bands in a manner different from that of 4A_2-5_ treated group. The cleavage preferences of Chimera-E-PO for these specific cleavage sites indicated its specific binding for RNA substrate.

### Evaluation of ASO-4A_2-5_ chimera for viral RNA knockdown and pseudovirus inhibition of SARS-CoV-2

We first selected envelope gene featured with a relative short viral RNA sequence, and evaluated E-RNA degradation efficiency using Chimera-E in Vero cells after co-transfection of pCAG-nCoV-E-FLAG plasmids. As we expected, treatment of 20 nM ASO-E alone could only partially downregulate E-RNA level to 83% in comparison to the negative control, while treatment of 20 nM Chimera-E downregulated E-RNA level to 35%, 2-fold more efficiently than that of ASO-E as measured by RT-qPCR (**Fig. 2A**). In addition, RNase L transcription level was also significantly increased with higher concentration of Chimera-E (**Fig. 2B**) which may further enhance the RNase L induced sequence-specific degradation of E-RNA. This result showed that Chimera-E could potently decreased intracellular E-RNA levels facilitated by RNase L activation, which inspired us to develop 4A_2-5_-ASO chimeras for spike protein, a more promising target to inhibit SARS-CoV-2 infection.

**Fig 2.**
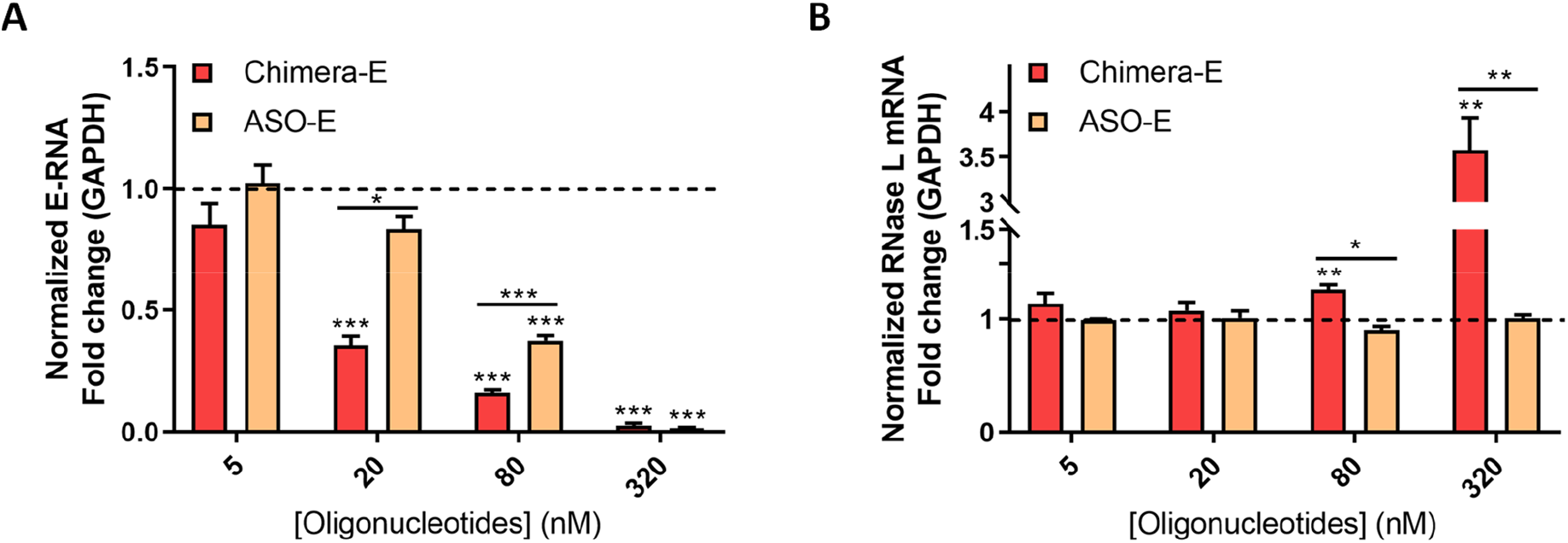
Targeted degradation of SARS-CoV-2 envelope RNA (E-RNA) with RNase L participation in Vero cells. Effective degradation of E-RNA **(A)** and up-regulation of RNase L **(B)** with the chimeric sequence targeting E-RNA of SARS-CoV-2 (Chimera-E) in comparison to pure antisense oligonucleotide (ASO-E) in Vero cells co-transfected with pCAG-nCoV-E-FLAG plasmids (250 ng/well), as measured by RT-qPCR. Data represent mean ± s.e.m. (n ≥ 3). *P < 0.033, **P < 0.002, ***P < 0.001 as measured by a two-tailed Student’s t test.

The on-target effects of three previously designed chimeric oligonucleotides (**Table S1**) against the spike RNA (S-RNA) of SARS-CoV-2 were evaluated in Vero cells using RT-qPCR (**Fig. 3A**). After co-transfection of pCAG-nCoV-S-FLAG plasmids and 80 nM oligonucleotides for 24 hours, all chimeras (Chimera-S) and antisense oligonucleotides (ASO-S) decreased S-RNA down to less than 50% level of negative control. Comparing ASO-S and Chimera-S containing the same antisense oligonucleotide sequence, more than 2-fold enhancement of S-RNA degradation was observed for Chimera-S that was able to activate endogenous RNase L by 4A_2-5_ moiety. Among three chimeric antisense oligonucleotides, Chimera-S4 displayed the highest enhancement in S-RNA degradation compared with its control group (ASO-S4). In addition, RNase L transcription levels in Vero cells upon the treatment of Chimera-S4, Chimera-S5 and Chimera-S6 were 2- ~ 4-fold higher than that of corresponding negative control, while all three antisense oligonucleotides without the conjugation of 4A_2-5_ had no obvious effects on RNase L transcription (**Fig. 3B**). These results reconfirmed that chimeric ASO conjugated with 4A_2-5_ efficiently activated cellular RNase L which further enhanced the degradation of target viral RNA.

**Fig 3.**
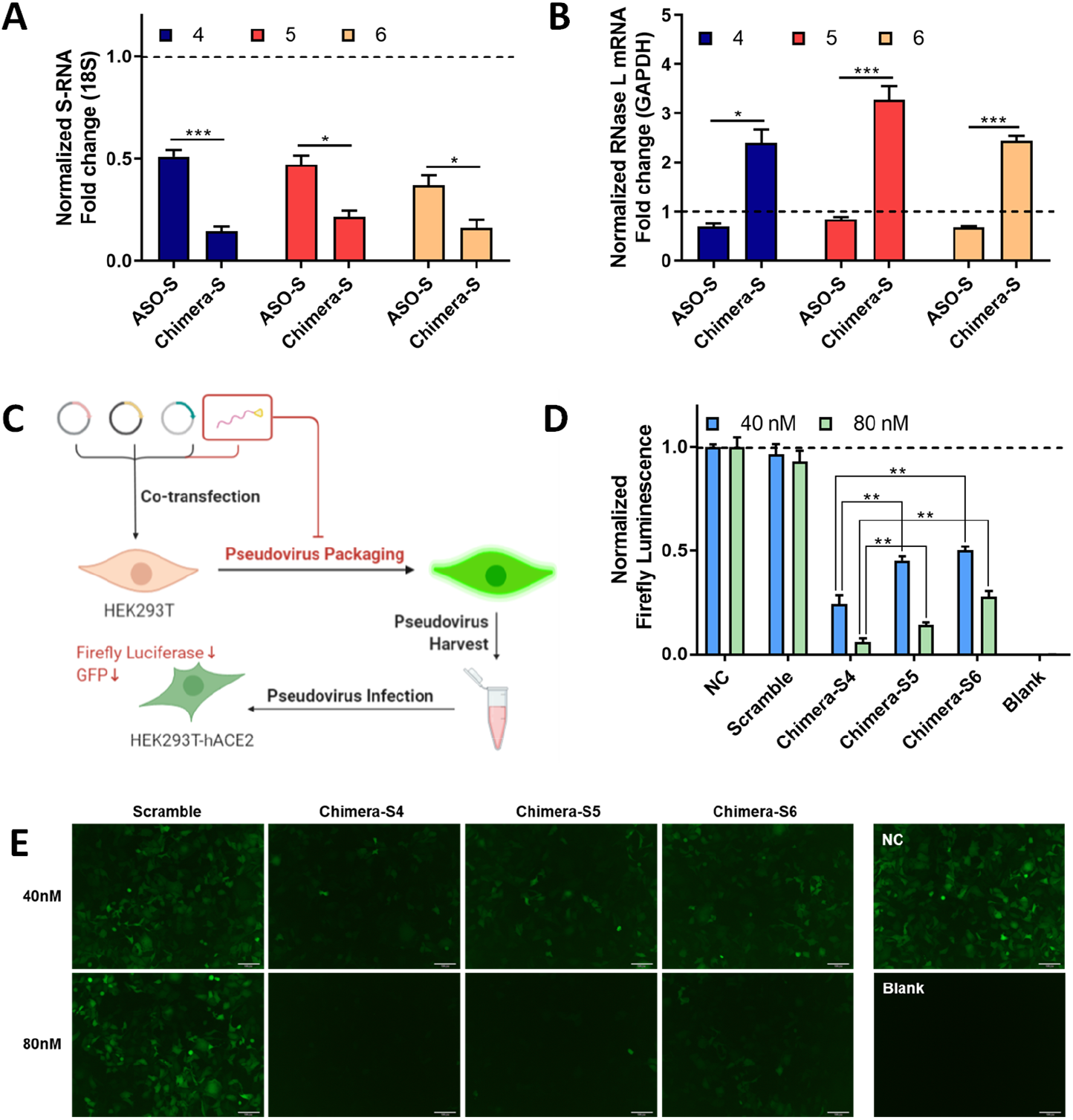
Screening the most effective 4A_2-5_-ASO chimeric oligonucleotides to target spike RNA (S-RNA) of SARS-CoV-2. Transcription levels of S-RNA **(A)** and RNase L mRNA **(B)** in 24h after co-transfection of pCAG-nCoV-S-FLAG plasmid and Chimera-S4, Chimera-S5, Chimera-S6 or their corresponding antisense oligonucleotides (ASO-S4, ASO-S5, ASO-S6) respectively, as measured by RT-qPCR. **(C)** Experimental procedure to evaluate the inhibitory effect of 4A_2-5_-ASO chimeras on virus packaging and infection in a pseudotyped SARS-CoV-2 infection model. Relative expression levels of firefly luciferase **(D)** and GFP **(E)** in infected HEK293T-hACE2 cells after Chimera-S4, Chimera-S5, or Chimera-S6 treatment (40 nM and 80 nM), respectively. Negative control (NC), group transfected with only virus-constructing plasmids. Scramble, group treated with the plasmids and a nonsense oligonucleotide Blank, group without exogenous transfection. Scale bar = 100 μm. Data represent mean ± s.e.m. (n ≥ 3). \**P* < 0.033, \*\**P* < 0.002, \*\*\**P* < 0.001 as measured by a two-tailed Student’s t test.

To further compare their efficiencies of S-RNA degradation and viral packaging inhibition, all above three chimeric oligonucleotides (Chimera-S4, Chimera-S5 and Chimera-S6) were applied to HEK293T packaging cells in a pseudotyped SARS-CoV-2 infection model (**Fig. 3C**). Since chimeric oligonucleotides led to the degradation of S-RNA, decrease of S protein expression and pseudovirus production could be observed. For titration of pseudovirus, two reporter genes, GFP and firefly luciferase were carried by pseudovirus. A hACE2 expressed cell line HEK293T-hACE2 was also established for pseudovirus titration. To examine effects of chimeric oligonucleotides on the transfection efficiency in pseudotyped SARS-CoV-2 infection model, GFP level in HEK293T packaging cell was compared. As expected, all groups showed similar GFP level (**Fig. S1**), indicating similar transfection efficiency in HEK293T packaging cells. In the infection model, Chimera-S4 was more potent to reduce titer of pseudovirus. At 40 nM and 80 nM, Chimera-S4 treatment reduced luminescence to 24% and 6% respectively, comparing to the corresponding control group, while firefly luminescence was down to 45% and 14% for Chimera-S5, 50% and 28% for Chimera-S6 at the same concentrations (**Fig. 3D**). Meanwhile, scrambled oligonucleotide showed no inhibition of pseudovirus, confirming the on-target effect of Chimera-S. GFP level of HEK293T-hACE2 was also monitored and Chimera-S4 also showed the most promising inhibition efficiency (**Fig. 3E**), which was consistent with luciferase assay. These results clearly showed that Chimera-S4 was the most effective and promising antiviral candidate among above Chimera-S oligonucleotides and could be used for further assessment.

### Chimera-S4 as a potent inhibitor of SARS-CoV-2 pseudovirus packaging

We further investigated the concentration dependence of Chimera S4 for S-RNA degradation. RT-qPCR results showed that 20 nM Chimera-S4 induced a reduction of S-RNA up to 80%. Increasing its concentration to 80 nM only led to slight enhancement of S-RNA reduction, but would cause an approximately 2-fold up-regulation of RNase L expression (**Fig. 4A, 4B**). Surprisingly, the titers of SARS-CoV-2 pseudovirus dropped sharply from 60% to 6% when the concentration of Chimera-S4 increased from 20 to 80 nM (**Fig. 4C**). In comparison to the individual ASO-S4 and 4A_2-5_, Chimera-S4 degraded S-RNA in Vero cells with up to 4.5- and 2.1-fold higher efficiency at 40 nM concentration (**Fig. 4A**). In addition, 2.9- and 1.4-fold higher upregulation of RNase L were also observed upon 80 nM Chimera S4 treatment (**Fig. 4B**). Compared with physically mixed 4A_2-5_ and ASO-S4 (4A_2-5_ + ASO-S4), Chimera-S4 led to similar reduction of S-RNA in Vero cells at 20 nM ~ 80 nM concentrations. However, the result of luciferase assays showed that Chimera-S4 displayed 3.8- and 19.2-fold higher inhibitory effects on viral titers at 40 nM and 80 nM than those of 4A_2-5_ + ASO-S4 group in HEK293T cells (**Fig. 4C**). To reconfirm the efficiency of Chimera-S4, GFP expression in infected HEK293T-hACE2 cells was also analyzed by flow cytometry. The positive rate of GFP fluorescent cells treated with 40 nM Chimera-S4 was 29.07%, much lower than those of negative control (81.46%), 4A_2-5_ (71.35%), ASO-S4 (69.06%) and 4A_2-5_+ASO-S4 (65.70%) groups under the same assay conditions (**Fig. 4D**), and could be further enhanced at higher concentrations of Chimera-S4 (**Fig. S2**). Fluorescent images of infected HEK293T-hACE2 cells were consistent with the results presented by flow cytometry (**Fig. 4E, Fig. S2**). Similarly, GFP fluorescence in HEK293T packaging cells confirmed the consistency of transfection efficiency across different groups (**Fig. S2**). All these results clearly showed that Chimera-S4 could efficiently reduce S-RNA level in a RNase L-facilitated manner and effectively inhibit SARS-CoV-2 pseudovirus at moderate doses, without serious damage to cell status or viability (**Fig. S2, Fig. S3**).

**Fig 4.**
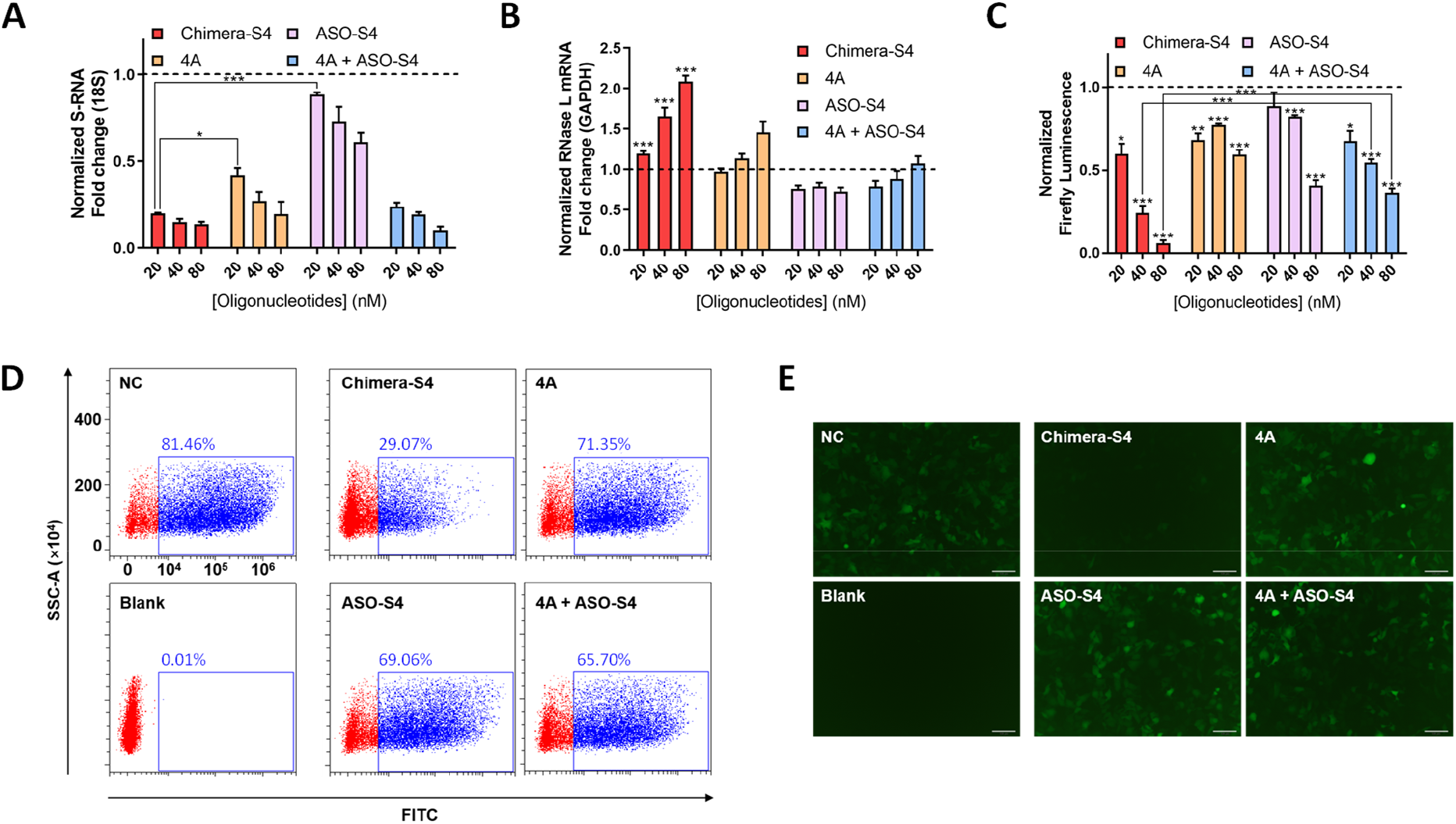
Chimeric-S4 can effectively degrade S-RNA of SARS-CoV-2 in Vero cells and inhibit pseudoviral infection of SARS-CoV-2 *in vitro.* **(A, B)** Concentration-dependent degradation of S-RNA and increase of RNase L mRNA in Vero cells after 24 h treatment with Chimera-S4. Firefly luminescence (**C,** 20 ~ 80 nM oligonucleotides), cytometry analysis of GFP signals (**D**, 40 nM oligonucleotides), GFP fluorescence images (**E**, 40 nM oligonucleotides) in HEK293T-hACE2 cells after 48 h infection of the collected SARS-CoV-2 pseudovirus under different treatments. Negative control (NC), transfection of virus-constructing plasmids. Blank, without exogenous transfection. 4A_2-5_ + ASO-S4, co-transfection of virus-constructing plasmids and physically mixed 4A_2-5_ and ASO-S4 with each final concentration of 20 nM, 40 nM and 80 nM. Scale bar = 100 μm. Data represent mean ± s.e.m. (n ≥ 3). \**P* < 0.033, \*\**P* < 0.002, \*\*\**P* < 0.001 as measured by a two-tailed Student’s t test.

As mentioned in the above results, Chimera-S4 obviously induced the up-regulation of RNase L, which also participated in antiviral immunity and activated the expression of other antiviral proteins, such as IFN-β and IL-6. Thus, we also assayed these intracellular mRNA levels of IFN-β and IL-6 after the activation of RNase L induced by Chimera-S4 (**Fig. S3**). When A549 cells were transfected with Chimera-S4 at different concentrations from 40 nM to 80 nM, the relative mRNA levels of IFN-β and IL-6 simultaneously increased from 5.9- to 26-fold and 2.0- to 7.5-fold in a concentration-dependent manner, respectively. In addition, Chimera-S4 induced much higher levels of antiviral proteins than those of individual 4A_2-5_, ASO-S4 and 4A_2-5_ + ASO-S4 mixture, indicating its higher potential to simultaneously activate antiviral immune response in SARS-CoV-2 therapy.

### Inhibition of SARS-CoV-2 pseudovirus mutants by Chimera-S4

Mutation ΔH69/ΔV70 and N501Y on spike protein of SARS-CoV-2 have been reported to cause S-gene target failure (SGTF) and greatly increase viral transmissibility (*20, 21*). To assess the broad-spectrum inhibition on SARS-CoV-2 mutants, Chimera-S4 was then co-transfected with pseudovirus packaging plasmids carrying ΔH69/ΔV70, N501Y or dual-site mutations into HEK293T packaging cells for further viral inhibition assay. Transfection efficiencies in HEK293T cells across different groups were consistent (**Fig. S4**). luciferase assay showed that titer of all three mutants were reduced to less than 20% after 48 hours treatment of 40 nM Chimera-S4 (**Fig. 5A**), which indicated a more robust inhibition of viral infection than those of 4A_2-5_, ASO-S4, 4A_2-5_ + ASO-S4 and scrambled sequences. GFP fluorescence analysis in infected HEK293T-hACE2 cells also displayed the same inhibiting manner as the firefly luciferase assay (**Fig. 5B**). These results indicated that Chimera-S4 could generally and efficiently inhibit the packaging and infection of ΔH69/ΔV70 and/or N501Y mutated SARS-CoV-2 pseudovirus *in vitro.*

**Fig 5.**
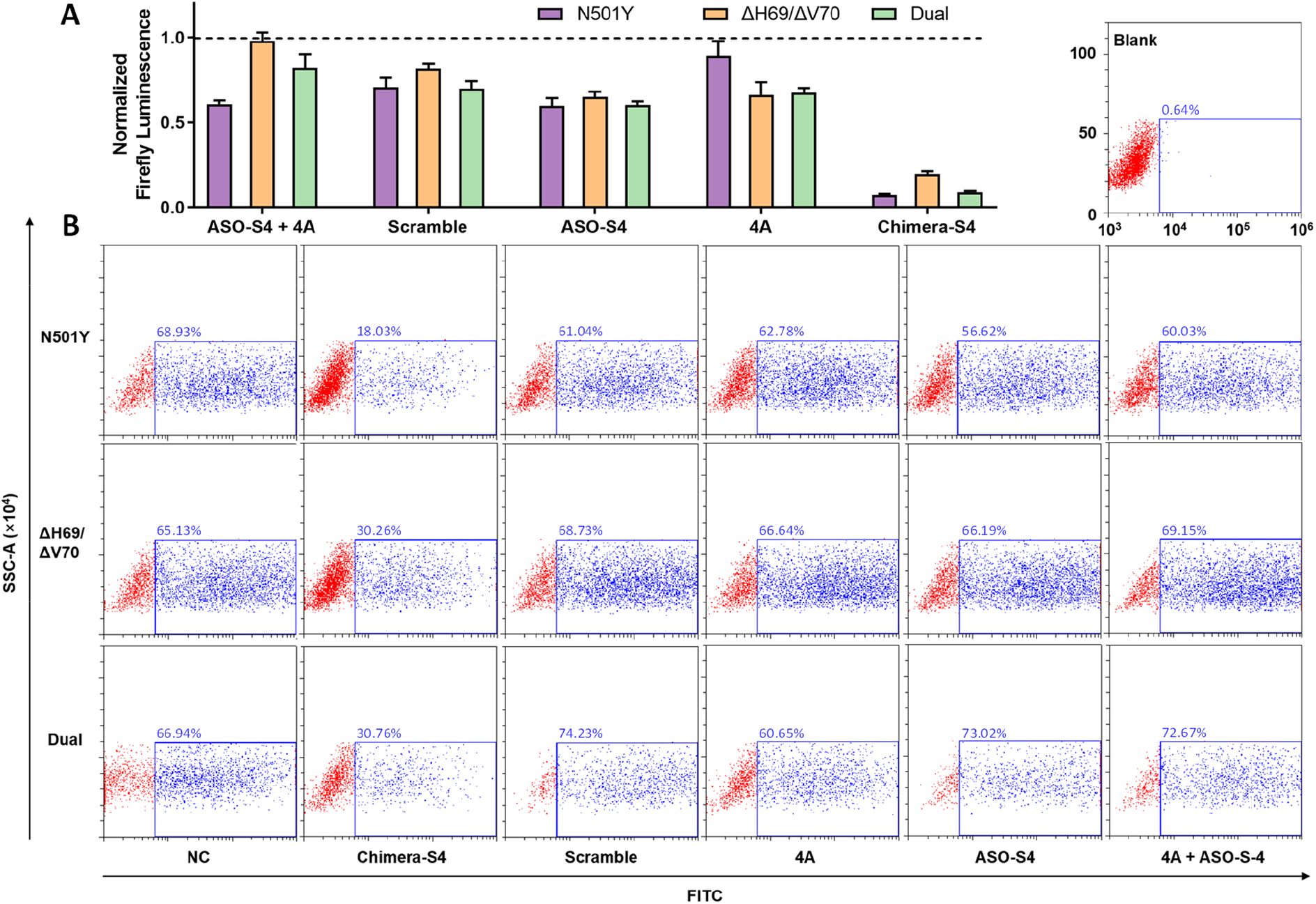
Investigate the viral titer of SARS-CoV-2 mutants upon Chimera-S4 treatment. Efficient inhibited infection of three mutated SARS-CoV-2 pseudoviruses, N501Y, ΔH69/ΔV70 and their combined mutants (Dual) in HEK293T-hACE2 cells after Chimera-S4 treatment (40 nM, 48 h), as measured by luciferase assay **(A)** and GFP signal analysis **(B)**.

## Discussion

We demonstrated that 4A_2-5_-chimeric antisense oligonucleotides (4A_2-5_-ASO) displayed potent antiviral effects against SARS-CoV-2 and its mutants by degrading target viral RNA of structural proteins through recruiting endogenous RNase L. *In vitro* cleavage assay validated that chimeric oligonucleotides could activate RNase L to cleave target RNA in a manner different from 4A_2-5_ induced substrate RNA cleavage, indicating its sequence-specific targeting process. Co-transfection of Chimera-E or Chimera-S with plasmids for E-RNA or S-RNA in Vero cells achieved up to 65% and 80% downregulation of RNA targets at 20 nM, which is much more efficient than that of corresponding classic ASOs, and much more unlikely to cause ubiquitous basal RNA decay due to 4A_2-5_ induced RNase L activation. Instead of IFN-deficient Vero cells, human alveolar basal epithelial cells A549 have been used to evaluate upregulation of IFN-β and IL-6 in a reprogrammed antiviral state upon RNase L activation (*9*). Upon treatment of Chimera-S4, the upregulation of antiviral gene transcription like IFN-β and IL-6 in A549 cells was observed, which is consistent with the intense activation of RNase L triggered by Chimera-S4. Due to the positive feedback of IFN-β on RNase L activation (*11*) and host defense against virus infection, 4A_2-5_-ASO chimera shows a strong inhibitory effect on virus proliferation at relatively low concentrations. Moreover, the mutation sites of spike genes corresponding to N501Y and ΔH69/ΔV70 are not overlaid with the target sequence of S-RBD RNA and can be still recognized by Chimera-S4, thus barely influencing its targeting process required for efficient S-RNA degradation. Therefore, in infection models of all three SARS-CoV-2 pseudovirus mutants, strong inhibition of SARS-CoV-2 packaging and infection was still successfully achieved.

Due to the acute shortage of biosafety level 4 conditions in the global COVID-19 pandemic, we currently have no chance to further evaluate the inhibitory effect of our chimeric 4A_2-5_-ASO on natural SARS-CoV-2 infection. However, we did confirm that chimeric oligonucleotides could efficiently inhibit viral proliferation in the SARS-CoV-2 pseudovirus infection model. And these exogenous ASO could be effectively delivered into lung tissues according to previously reported delivery strategies (*22, 23*), which would further expand their potential applications *in vivo.* Furthermore, interferons themselves are also therapeutic agents used for virus immunotherapy (*24, 25*) if severe release of proinflammatory cytokines is well controlled.(*26*).

In conclusion, we developed a group of 4A_2-5_ chimeric oligonucleotides based on nucleic acid-hydrolysis targeting chimera strategy (NATAC) and successfully down-regulated target SARS-CoV-2 RNAs. Among them, Chimera-S4 showed the most potent degradation of S-RBD RNA and the inhibition of SARS-CoV-2 pseudovirus. Compared with classic ASO silencing strategy, Chimera-S4 also activated RNase L, which significantly improved RNA degradation efficiency, and induced additional antiviral immune response upon its recruitment by 4A_2-5_ ligand. This chimeric sequence still showed robust inhibiting capability toward three highly transmissible SARS-CoV-2 mutants involving N501Y and ΔH69/ΔV70 mutations. Antisense oligonucleotides have the characteristics of sequence-specific targeting, convenient design and synthesis, which make this 4A_2-5_-ASO chimera suitable for further development of nucleic acid drugs combating foreseeable evolving COVID-19 pandemics.

## Materials and Methods

### Experimental Design

We developed chimeric antisense oligonucleotides with enhanced degradation of target viral RNA and potent antiviral efficiency against SARS-CoV-2 facilitated by RNase L. Sequences of oligonucleotides, protein of purified GST-RNase L, plasmids and cell lines required by pseudotyped SARS-CoV-2 infection model were prepared first. Then *in vitro* RNase L cleavage assay and primary RT-qPCR assay in Vero cells were carried out to validate the RNase L recruiting mechanism of the chimeric design. Next, we screened three chimera candidates targeting S-RBD gene of SARS-CoV-2 for the most efficient one through evaluating their downregulation of S-RNA by RT-qPCR in Vero cells and the inhibition of SARS-CoV-2 pseudovirus packaging in HEK293T cells. Mutants of SARS-CoV-2 pseudovirus involving N501Y and/or ΔH69/ΔV70 mutations were also included. Further concentration dependence, regulation of IFN-β and IL-6 as well as influence on cell viability of the most efficient oligonucleotide candidate were investigated in Vero cells or A549 cells.

### Design of oligonucleotide sequences

Secondary structures of SARS-CoV-2-E RNA and SARS-CoV-2-S-RBD RNA were predicted by RNAfold web server (Institute for Theoretical Chemistry, University of Vienna) based on minimum free energy (MFE) and partition function algorithms. Antisense oligonucleotide candidates targeting specific SARS-CoV-2 RNA fragments were given by Oligowalk (Mathews group, University of Rochester Medical Center) for the further selection of antisense oligonucleotides.

### Preparation of oligonucleotides

Chimeric oligonucleotides (Chimera-E or Chimera-S), ASO-S control oligonucleotides and 3’-Cy3 labeled E-RNA segment were purchased from Biosyntech. Chi-E-PO, ASO-E and 4A_2-5_ control oligonucleotide were synthesized on ABI DNA/RNA synthesizer based on standard phosphoramidite chemistry, and were purified through HPLC (Waters, Alliance e2695) after the cleavage and deprotection. All the oligonucleotides were confirmed by ESI-MS (Sangon Biotech). Each oligonucleotide was dissolved in nuclease-free water and quantified with NanoDrop 2000 (Thermo Fisher Scientific) at 260 nm before use.

### Preparation of plasmids

The pCAG-FLAG vectors containing SARS-CoV-2-E gene (pCAG-nCoV-E-FLAG) or SARS-CoV-2-S gene (pCAG-nCoV-S-FLAG) were generously provided by Prof. Wang Pei-Hui’s lab (Shandong University). Full length RNase L gene was synthesized and subcloned into pGEX-4T-3 vector (pGEX-4T-RNaseL-GST) by GENEWIZ as previously described (*27*).

Plasmid pcDNA 3.1-SARS-CoV-2-Spike, pLVX-hACE2-IRES-puro, pMD2G-VSVG, pspAX.2, pLenti-FLuc-GFP were constructed to generate SARS-CoV-2 pseudovirus and establish transgenic cell line HEK293T-hACE2. Briefly, gene segment containing spike protein of SARS-CoV-2 wa s synthesized by GenScript Inc. without codon optimization and was inserted into pcDNA 3.1 to g et pcDNA 3.1-SARS-CoV-2-Spike using NEBuilder® HiFi DNA Assembly Master Mix (NEB) according to the manufacturer’s instructions. In order to construct transfer plasmid pLVX-hACE2-IRES-puro, plasmid containing complete ORF of hACE29 (pMD18-T-hACE2) was purchased fro m Sino biological Inc. and hACE2 gene was sequenced by BGI Inc. Then hACE2 segment was amplified by primer F 5’-ATGTCAAGCTCTTCCTGG-3’ and primer R 5’-CTAAAAGGAGGTC TGAACATC-3’, then restriction enzyme cutting site XhoI and XbaI was added using primer forw ard: 5’-CTCGAGCTCGAGGCCGCCACCATGTCAAGCTCTTCCTGGC-3’ and reverse: 5’-TC TAGATCTAGACTAAAAGGAGGTCTGAACATCA-3’. Lentiviral transfer plasmid pLVX-IRE S-puro was stored in our lab. Insertion of hACE2 into pLVX-IRES-puro was conducted by double digestion of XhoI and XbaI (Fermantas) and ligation of T4 ligase (NEB) according to manufacturer’s instructions. Plasmid pMD2G-VSVG, pspAX.2, pLenti-FLuc-GFP was stored in our lab (*28*).

Mutant plasmids pCMV-hnCoV-S-H501Y (forward: 5’-CCAGCCTACATATGGCGTGGGCT-3’, reverse: 5’-AAGCCGTAAGACTGGAGTG-3’) and pCMV-hnCoV-S-Δ69/70 (forward: 5’-TCCGGCACAAACGGCACA-3’, reverse: 5’-GATGGCGTGGAACCATGTC-3’) were obtained from the wild type plasmids pCMV-hnCoV-S via Q5 SiteDirected Mutagenesis Kit (NEB). pCMV-hnCoV-S-H501Y-Δ69/70 was obtained from pCMV-7.1-hnCoV-S-H501Y (forward: 5’-TCCGGCACAAACGGCACA-3’, reverse: 5’-GATGGCGTGGAACCATGTC-3’) via Q5 SiteDirected Mutagenesis Kit (NEB). All plasmids were confirmed by gene sequencing (BGI Beijing). All plasmids used for transfection were amplified using a Maxiprep kit (Promega), according to the manufacturer’s instructions.

### Preparation of RNase L-GST protein

The RNase L-GST fusion protein was expressed in *Escherichia coli* strain DH5α transformed with pGEX-4T-RNaseL-GST plasmid as previously described (*27*). Briefly, cells were grown at 30 C to A595 = 0.5, then 0.1 mM isopropylthio-galactoside was added and cells were grown for another 3 h at 30 °C before harvest. After centrifugation at 4000 rpm at 4 °C for 15 min, cells were washed with 0.8% NaCl and resuspended in 50 mL buffer A (10 mM NaH2PO4, pH7.4, 600 mM NaCl, 10% glycerol, 1 mM EDTA, 0.1 mM ATP, 5 mM MgCl2, 14 mM 2-mercaptoethanol, 1 μg/mL leupeptin) supplemented with 1% Triton X-100, 1 mM PMSF, 1 μg/mL lysozyme and 10 mM DTT. Then cells were sonicated on ice and cell lysates were centrifugated at 11, 000 rpm at 4 °C for 40 min to collect supernatants. RNase L-GST protein in supernatants was purified via GST affinity chromatography (HP, Cytiva) with buffer B (20 mM glutathione, 300 mM NaCl, 50 mM Tris-HCl, pH 8.0, 1 μg/mL leupeptin) as eluent. Fractions containing RNase L-GST protein were collected and the purity of the protein were analyzed by SDS-polyacrylamide gel electrophoresis.

### *In vitro* Cleavage of E-mRNA by RNase L

Conditions for RNase L cleavage of single strand RNA were formerly reported (*13*). Briefly, Cy3-labeled E-RNA fragment as the substrate RNA was folded in 1× RNase L NM Buffer (25 mM Tris-HCl, pH7.4, 100 mM KCl) at 8 μM by heating the solution at 95 °C for 30 s and slowly cooling to 25 °C. Then the above solution was supplemented with 2× Supplementary Buffer (25 mM Tris-HCl, pH7.4, 100 mM KCl, 20 mM MgCl2, 14 mM β-mercaptoethanol, 100 μM ATP) and aliquots of 4A_2-5_ or Chi-E-PO was then added, followed by incubation at 25 °C for 30 min. Both 4A_2-5_ and Chimera-O-E were diluted in 1× RNase L M Buffer (25 mM Tris-HCl, pH 7.4, 100 mM KCl, 10 mM MgCl2, 7 mM β-mercaptoethanol, 50 μM ATP). Then RNase L was added at an equimolar concentration of 4A_2-5_ or Chimera-O-E. Each sample was supplement to a final volume of 8 μL and was further incubated at 25 °C for 60 min. After quenching RNase L cleavage by adding 2× Loading Buffer (8 M urea, 2 mM Tris-base, 20 mM EDTA, 0.01% bromophenol blue and 0.01% xylene cyanol), samples were heated at 95 °C for 3 min and loaded in a denaturing 12.5% polyacrylamide gel. The gel was run at 250 V for 20 min and imaged using Chemiluminescence gel imaging system (ChemiDoc XRS).

### Cell culture and Transfection Procedure

Vero cells and A549 cells were grown at 37 °C, 5% CO_2_ in DMEM (M&C) supplemented with 10% fetal bovine serum (PAN), 100 units/mL penicillin, and 100 μg/mL streptomycin. Cells were seeded and incubated for 24 h. Transfection of oligonucleotides and/or plasmids were performed using Lipofectamine™ 2000 (Invitrogen) according to the manufacturer’s instructions. After 6 h incubation, cells were replaced with fresh medium and incubated at 37°C, 5% CO_2_ in for another 18 h.

HEK293T cells and transgenic cell line HEK293T-hACE2 were maintained in DMEM (Gibco) supplemented with 10% fetal bovine serum (Gibco), 100 units/mL penicillin, and 100 μg/mL streptomycin. HEK293T cells were stored in our lab (*29*). Transfection of oligonucleotides and plasmids in pseudovirus infection models were performed using Lipofectamine™ 3000 (Invitrogen) according to the manufacturer’s instructions. After 6 h incubation, cells were replaced with fresh medium and incubated at 37°C, 5% CO_2_ in for another 42 h.

### Real-time Polymerase Chain Reaction

Vero cells or A549 cells were seeded into 24-well plates with the density of 7.5×10^4^ cells per well (for A549 cells, the density is 1×10^5^ cells per well). Oligonucleotides and/or plasmids (250 ng per well) were transfected to each well according to the group setting. After 24 h incubation at 37 °C, total RNA was extracted using BioZol reagent (Bioer) according to the manufacturer’s instructions. cDNAs were synthesized with HiScript III 1^st^ cDNA Synthesis Kit (+gDNA wiper) (Vazyme Biotech). Real time-polymerase chain reactions were performed with GoTaq qPCR Master Mix (Promega) according to the manufacturer’s instructions and completed on QuantStudio 6 Flex system (ABI). RNA expression levels were determined through the ΔΔCt method and normalized with GAPDH or 18S as a housekeeping gene.

### SRB Assay

Oligonucleotides and plasmids were transfected into Vero cells (seeded in 96-well plates with the density of 2×10^4^ cells per well). Cells were replaced with fresh medium after 6 h incubation. After another 18 h, culture medium was removed and cold 10% TCA was added (100 μL per well). The plate was incubated at 4 °C for 1 h and then washed with deionized water (200 μL per well) for four times. After naturally drying, 4 mg/mL Sulforhadamine B (SRB) dissolved in 1% aqueous acetic acid was added (100 μL per well) and the plate was incubated at room temperature for 30 min. Each well was rinsed with 1% acetic acid for five times and naturally dried. Finally, 10 mM unbuffered Tris base (pH 10.5) was added (100 μL per well). Read the optical density at 540 nm by a microplate reader (SYNERGY H1, BioTek).

### Establishment of transgenic cell line HEK293T-hACE2

Procedure to establish a cell line expressing human angiotensin-converting enzyme 2 (hACE2) receptor was previously described (*30*) and introduced in brief. HEK293T cells were used for lentiviral vector packaging and transduction. The cells were cultured in DMEM supplemented with 10% FBS (Gibco) and 1 mM nonessential amino acids (Gibco). Sub confluent HEK293T cells in 6-well plates were co-transfected with 0.72 μg of pLVX-hACE2-IRES-puro transfer plasmid, 0.64 μg of pMD2G-VSVG and 0.64 μg of pspAX.2 using transfecting reagent Megatran 1.0 (Origene).

Then, 6 h post transfection, the medium was replaced by DMEM supplemented with 3% FBS and 1 mM nonessential amino acids. Next, the lentiviral-containing supernatant was harvested at 48 h post transfection and filtered by a 0.45 μm filter (Pall). The resultant lentiviruses were used to integrate hACE2 gene into the genome of HEK293T cells. Procedure of stable lentiviral transduction was carried out as follows: HEK293T cells were seeded in a 6-well plate and transducted 24 h later with lentiviral filtrate in presence of 8 μg/mL polybrene (Macgene). Then, selection was performed under the pressure of 1 μg/mL puromycin (Invitrogen) until cells died completely. Then the cell line was verified by western blot.

### Generation of SARS-CoV-2 pseudovirus

Construction of a VSV pseudovirus carrying the spike protein of SARS-CoV-2 was formerly reported (*30*) and introduced in brief. HEK293T cells were used for pseudovirus packaging. Subconfluent HEK293T cells in 6-well plates were co-transfected with 1.2 μg of pLenti-FLuc-GFP transfer plasmid, 0.4 μg of pcDNA 3.1-SARS-CoV-2-Spike plasmid, 0.4 μg of pspAX.2 plasmid and oligonucleotides (0.3~1.1 μg) per well. 6 h post transfection, the medium was replaced by DMEM supplemented with 10% FBS. Next, cell status and green fluorescence was captured by inverted fluorescence microscope (Olympus) 48 h post transfection, then the pseudovirus-containing supernatant was harvested and filtered by a 0.45 μm filter (Pall). The resultant pseudoviruses were further analyzed for viral tilter by flow cytometry and luciferase assay.

### Pseudovirus infection and luciferase assay

In order to determine the titration of pseudovirus, expression of firefly luciferase was conducted as follows: HEK293T-hACE2 cells were seeded into 96-well black/clear bottom plates (Nunc) at 5×10^3^ cells per well and cultured for 24 h. Then the medium was replaced by 100 μL pseudovirus pLenti-FLuc-GFP filtrate and cells were incubated for another 48 h. Expression of firefly luciferase was quantitated by Bright Glo™ luciferase assay system (Promega) and the plates were read using a plate reader (Tecan Infinite M2000 PRO).

### Flow cytometry

The transfection efficiency during pseudovirus packaging was analyzed by flow cytometry. Briefly, HEK293T cells transfected with plasmid pcDNA pLenti-FLuc-GFP, pcDNA 3.1-SARS-CoV-2-Spike, pspAX.2 and oligonucleotides were incubated for 48 h and GFP expression level was analyzed by CytoFLEX flow cytometer (Beckman).

In order to confirm the titration of pseudovirus, HEK293T-hACE2 cells were seeded into 6-well plates. After 24 h incubation, the medium was replaced by 1 mL fresh medium mixed with 1 mL pseudovirus pLenti-FLuc-GFP filtrate. Cells were incubated for 48 h and GFP expression level was analyzed by CytoFLEX flow cytometer (Beckman).

### Statistical Analysis

GraphPad Prism 7.04 was used for statistical analysis and graphing. Two-tailed Student’s t test was used to compare data of two experimental groups.

## Author contributions

X.T. and X.S. conceived this study and designed experiments. X.S. and Q.W. prepared the protein. B.C. and Z.G. constructed pseudovirus mutants. W.M. and B.C. performed experiments related to pseudovirus. X.S. performed most of the experiments and analyzed data except those noted. X.S wrote the manuscript. X.T., Q.W., W.M. and D.Z. revised the manuscript.

## Data and materials availability

Genome sequences and protein sequence have been deposited in GenBank with accession number NC_045512.2, NM_021133.4 and NP_066956.1. All data needed to evaluate the conclusions in the paper are present in the paper and/or the Supplementary Materials. Additional data related to this paper are available from the corresponding author.

