## supplemental Figure S1-6 and Supplemental Table S1-2 for "Efficient Inhibition of SARS-CoV-2 Using Chimeric Antisense Oligonucleotides through RNase L Activation"

**This PDF file includes:**

Figs. S1 to S6
Tables S1 to S2

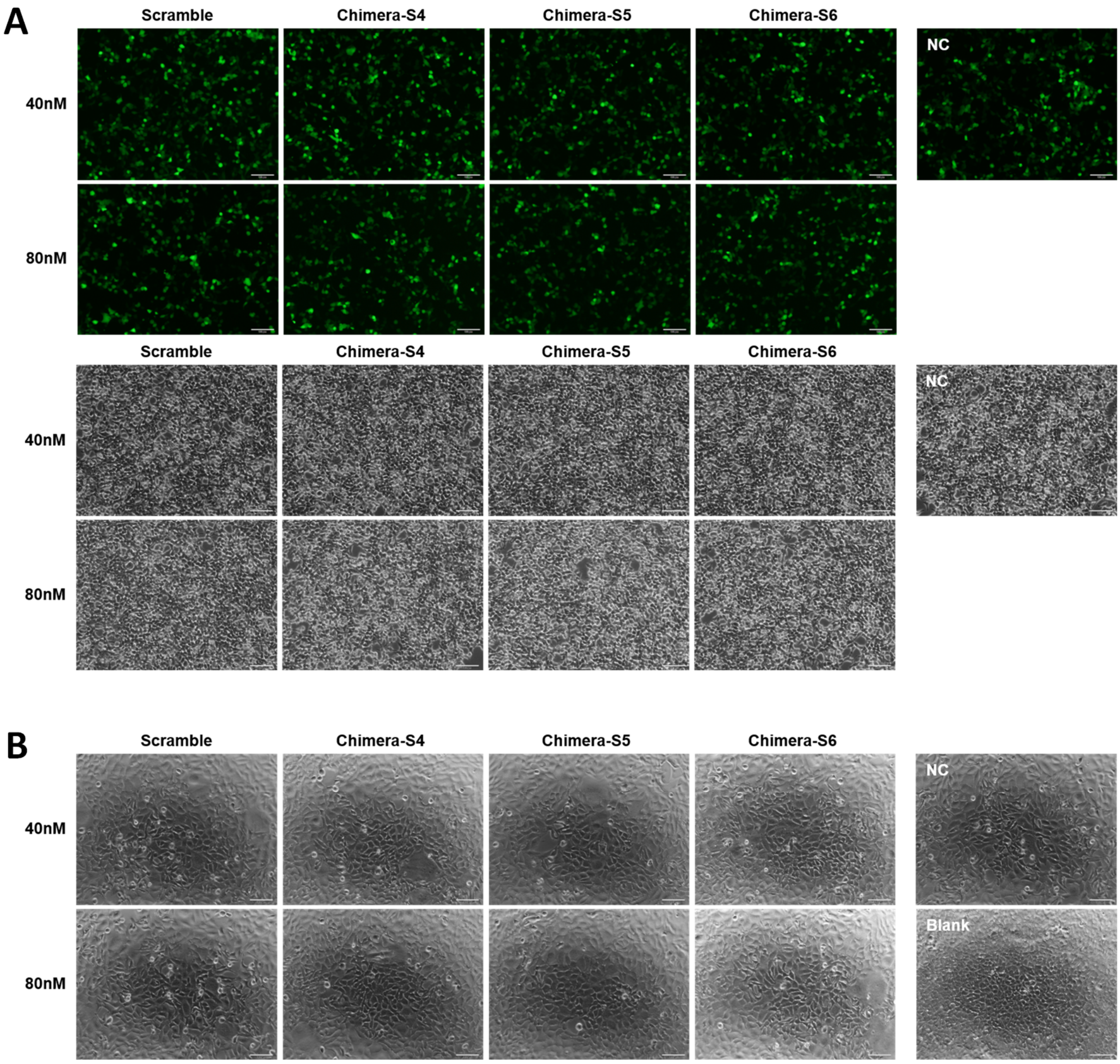


**Fig. S1.** Transfection efficiency and cell status in pseudotyped SARS-CoV-2 model during Chimera-S screening stage. **(A)** Fluorescent images and brightfield images of HEK293T cells incubated for 48 h after transfection of 40 nM or 80 nM Chimera-S4, Chimera-S5, Chimera-S6 or scrambled oligonucleotides. Similar green fluorescence of GFP between experimental groups and negative control (NC) group showed nearly equal transfection efficiency. **(B)** Brightfield images of HEK293T-hACE2 cells incubated for 48 h after infection. Scale bar = 100 µm.

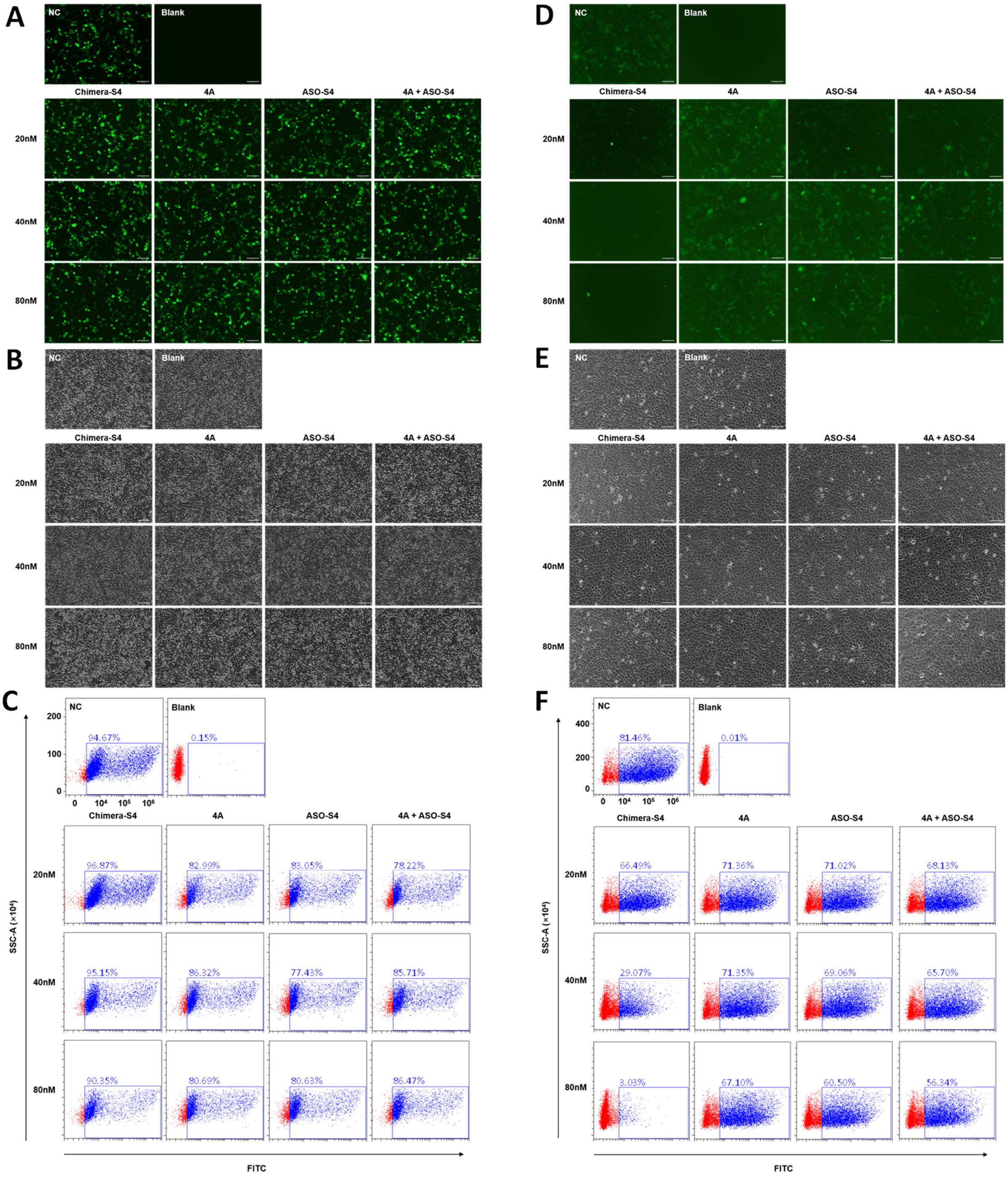


**Fig. S2.** Chimera-S4 inhibited assembly of SARS-CoV-2 in concentration dependent manner. Fluorescent images **(A)**, brightfield images **(B)** and quantified GFP signals **(C)** of HEK293T cells incubated for 48 h after transfection of different concentrations of Chimera-S4, 4A, ASO-S4 and 4A + ASO-S4. For HEK293T-hACE2 cells post 48 h infection, fluorescent images **(D)**, brightfield images **(E)** were also captured and their GFP signals **(F)** were analyzed by flow cytometry.

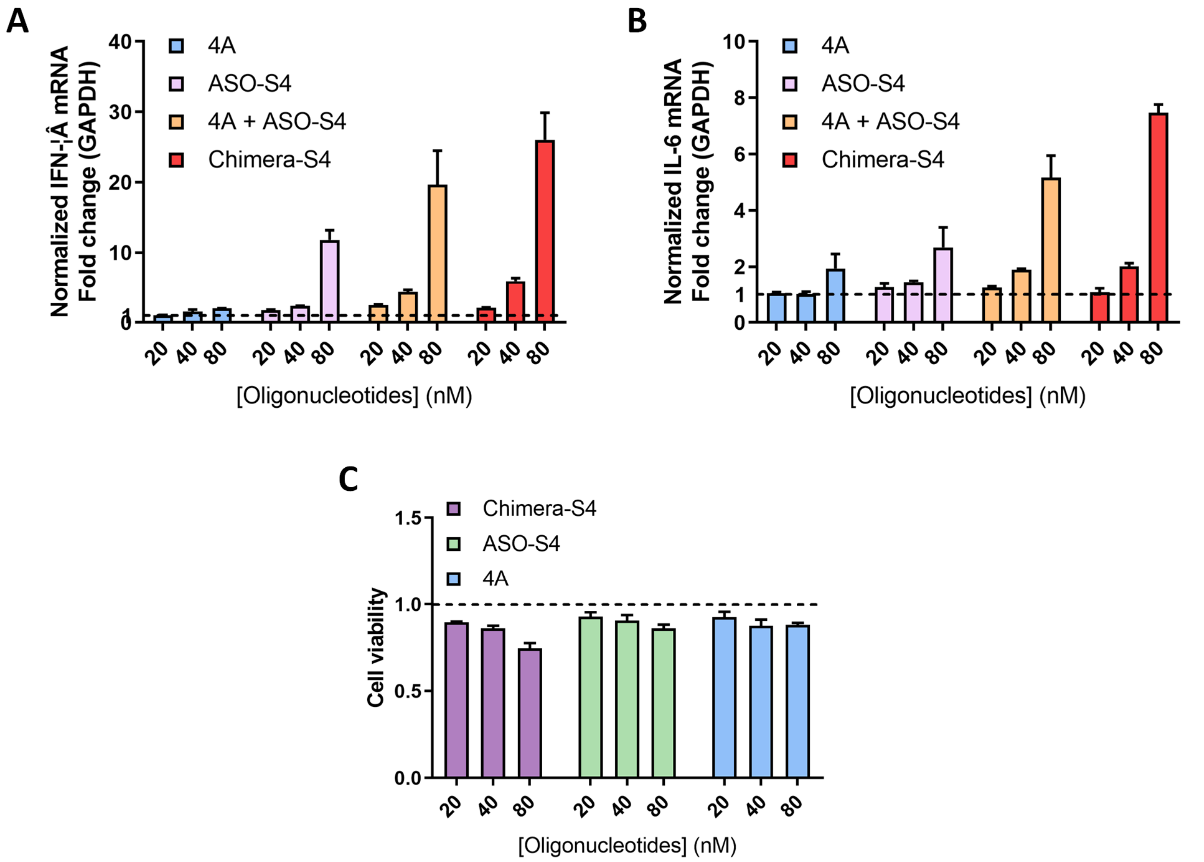


**Fig. S3.** Intracellular mRNA levels of IFN-β **(A)** and IL-6 **(B)** in Vero cells incubated for 24 h after transfection of different concentrations of Chimera-S4, 4A, ASO-S4 and 4A + ASO-S4, as measured by RT-qPCR. **(C)** Cell cytotoxicity assay in Vero cells. Data represent mean ± s.e.m. (n ≥ 3).

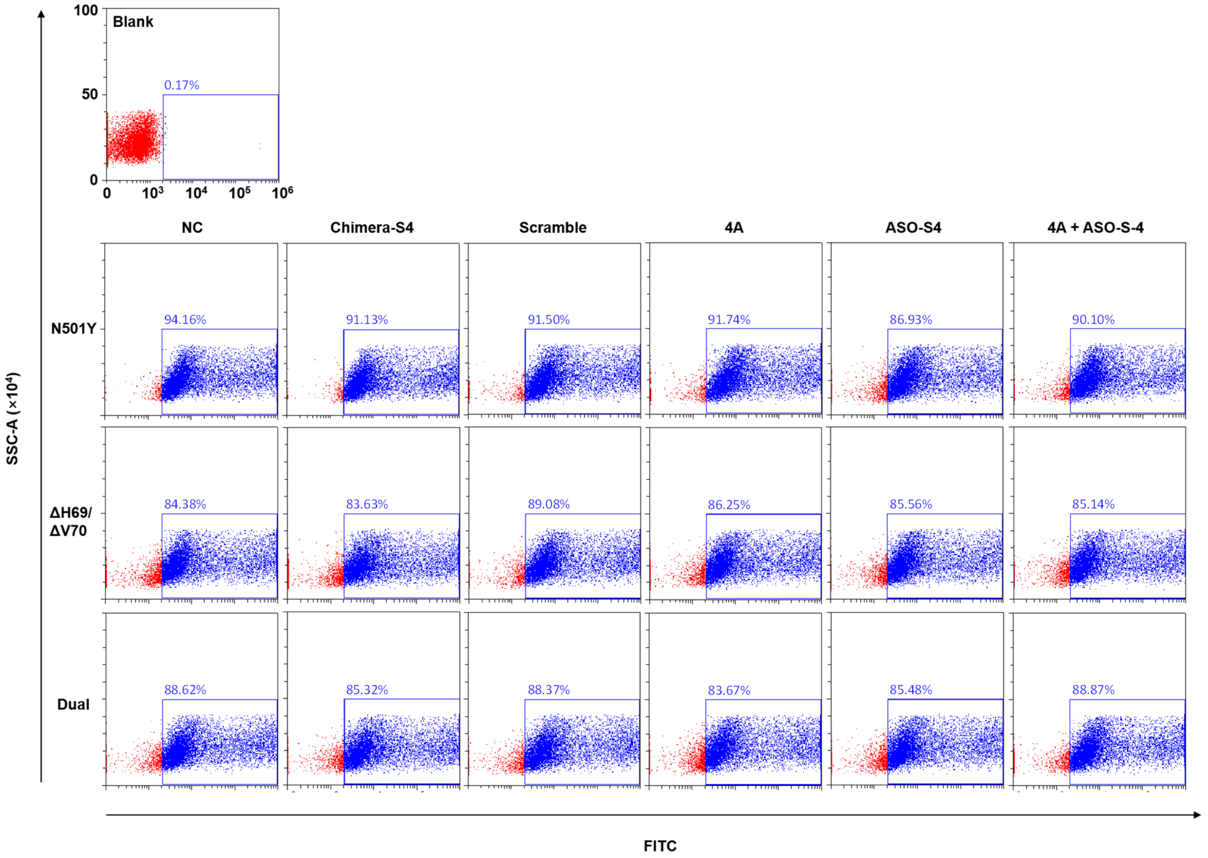


**Fig. S4.** Flow cytometry analysis of GFP expression in HEK293T cells packaging mutated SARS-CoV-2 pseudovirus, N501Y, ΔH60/ΔV70 or their combined mutants (Dual), for 48 h.

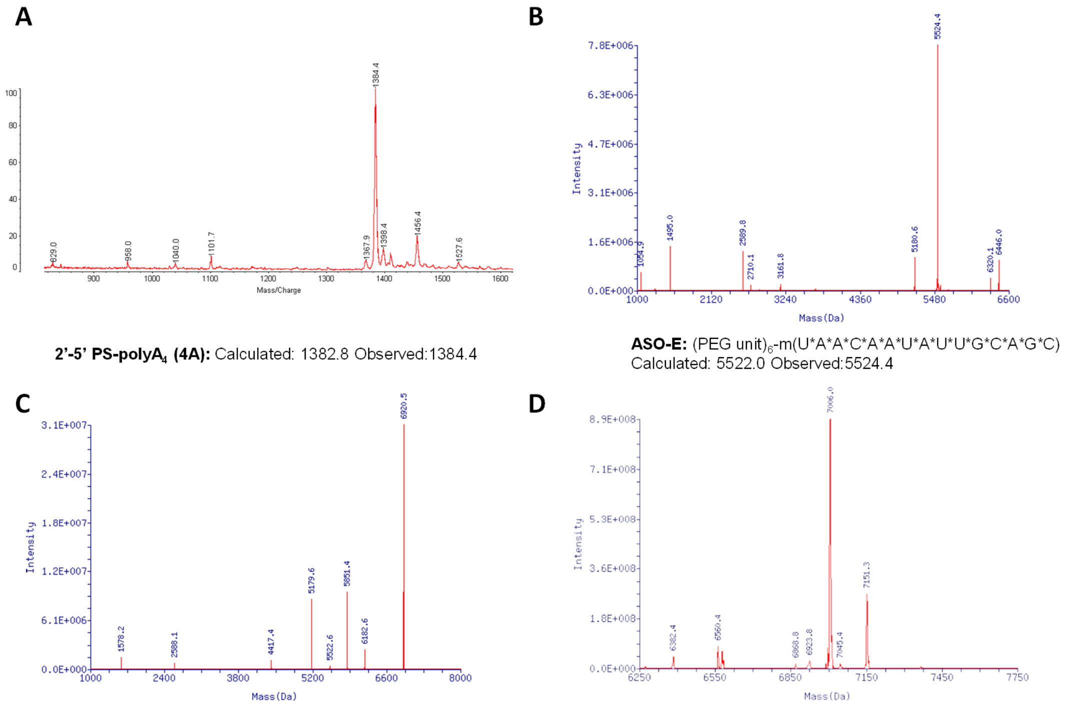


**2'-5' PS-polyA$_4$ (4A):** Calculated: 1382.8 Observed:1384.4

**ASO-E:** (PEG unit)$_6$-m(U*A*A*C*A*A*U*A*U*U*G*C*A*G*C)
Calculated: 5522.0 Observed:5524.4

**Chi-E-PO:** p-r(XXXX)-p-(PEG unit)$_6$-p*-m(U*A*A*C*A*A*U*A*U*U*G*C*A*G*C)
Calculated: 6918.9 Observed:6920.5

**Chimera-E:** p*-r(X*X*X*X)-p*-(PEG unit)$_6$-p*-m(U*A*A*C*A*A*U*A*U*U*G*C*A*G*C)
Calculated:7015.7 Observed:7006.0

**Fig. S5.** Mass spectra of 2'-5' polyA4 (**A**), ASO-E (**B**), Chi-E-PO (**C**) and Chimera-E (**D**). X = 2'-5'linked adenine nucleotide; *: phosphorothioate; m: 2'-*O*-methyl.

A

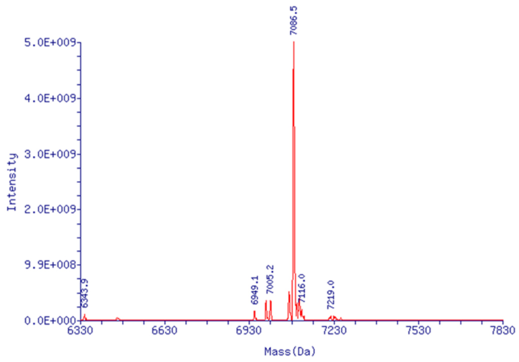


**Chimera-S4:** p*-r(X*X*X*X)-p*-(PEG unit)$_6$-p*-r(U*A*U*G*A*U*U*G*U*A*A*A*G*G*A)
Calculated:7081.7 Observed: 7086.5

B

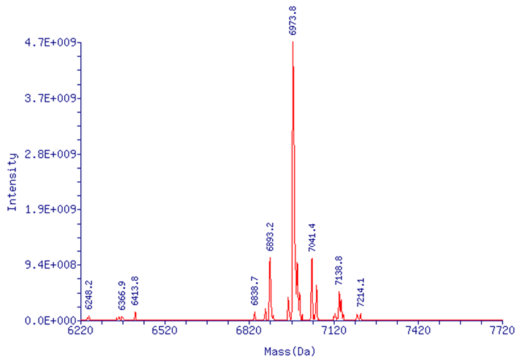


**Chimera-S5:** p*-r(X*X*X*X)-p*-(PEG unit)$_6$-p*-r(A*C*U*A*C*U*C*U*G*U*A*U*G*G*U)
Calculated:6970.5 Observed: 6973.8

C

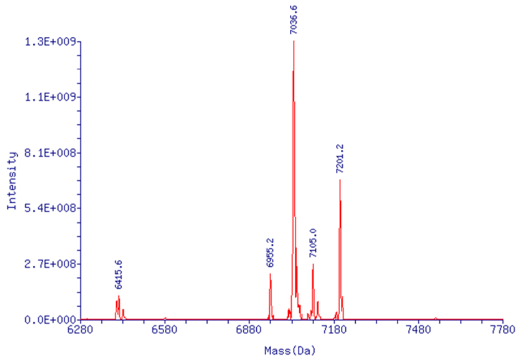


**Chimera-S6:** p*-r(X*X*X*X)-p*-(PEG unit)$_6$-p*-r(A*U*C*A*G*U*A*G*U*G*U*C*A*G*C)
Calculated:7032.6 Observed: 7036.6

D

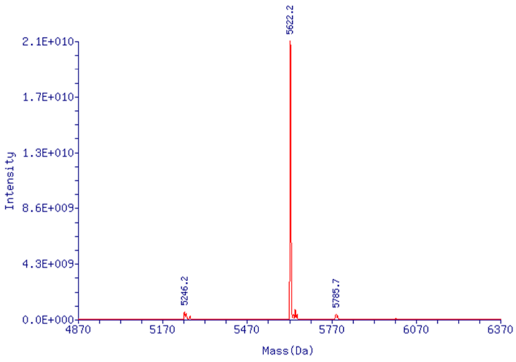


**ASO-S4:** (PEG unit)$_6$-p*-r(U*A*U*G*A*U*U*G*U*A*A*A*G*G*A)
Calculated:5620.6 Observed: 5622.2

E

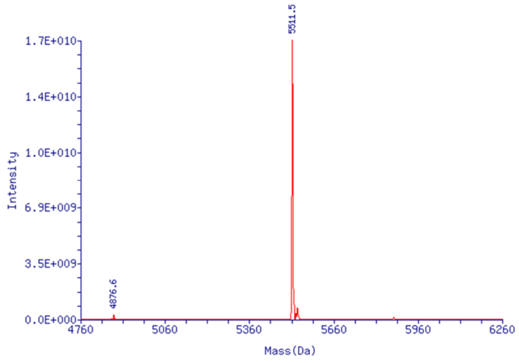


**ASO-S5:** (PEG unit)$_6$-p*-r(A*C*U*A*C*U*C*U*G*U*A*U*G*G*U)
Calculated:5509.5 Observed: 5511.5

F

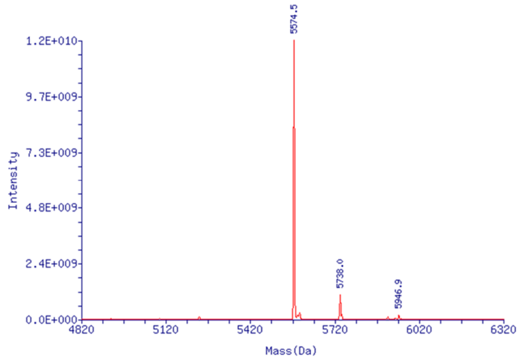


**ASO-S6:** (PEG unit)$_6$-p*-r(A*U*C*A*G*U*A*G*U*G*U*C*A*G*C)
Calculated:5571.5 Observed: 5574.5

**Fig. S6.** Mass spectra of Chimera-S4 (**A**), Chimera-S5 (**B**), Chimera-S6 (**C**), ASO-S4 (**D**), ASO-S5 (**E**) and ASO-S6 (**F**). X = 2’-5’linked adenine nucleotide; *: phosphorothioate; m: 2’-*O*-methyl.

**Table S1.** Oligonucleotides synthesized in this study.

| Name | Sequence (5'→3') |
|---|---|
| Chimera-S4 | p*-r(X*X*X*X)-p*-(PEG unit)$_6$-p*-m(U*A*U*G*A*U*U*G*U*A*A*A*G*G*A) |
| Chimera-S5 | p*-r(X*X*X*X)-p*-(PEG unit)$_6$-p*-m(A*C*U*A*C*U*C*U*G*U*A*U*G*G*U) |
| Chimera-S6 | p*-r(X*X*X*X)-p*-(PEG unit)$_6$-p*-m(A*U*C*A*G*U*A*G*U*G*U*C*A*G*C) |
| ASO-S4 | (PEG unit)$_6$-p*-m(U*A*U*G*A*U*U*G*U*A*A*A*G*G*A) |
| ASO-S5 | (PEG unit)$_6$-p*-m(A*C*U*A*C*U*C*U*G*U*A*U*G*G*U) |
| ASO-S6 | p*-(PEG unit)$_6$-p*-m(A*U*C*A*G*U*A*G*U*G*U*C*A*G*C) |
| Chimera-E | p*-r(X*X*X*X)-p*-(PEG unit)$_6$-p*-m(U*A*A*C*A*A*U*A*U*U*G*C*A*G*C) |
| Chi-E-PO | p-XXXX-p-(PEG unit)$_6$-p*-m(U*A*A*C*A*A*U*A*U*U*G*C*A*G*C) |
| ASO-E | (PEG unit)$_6$-p*-m(U*A*A*C*A*A*U*A*U*U*G*C*A*G*C) |
| 3'-Cy3 E-RNA | ACUGCUGCAAUAUUGUUAACGUGAGUCUUGUAAAACCUUCUUUUUACGUUUACUCUCG UGUU-Cy3 |

X = 2'-5'linked adenine nucleotide
*: Phosphorothioate
m: 2'-*O*-methyl

**Table S2.** Primers used for RT-qPCR.

| Primer | Sequence (5'→3') |
|---|---|
| GAPDH F | TGCACCACCAACTGCTTAGC |
| GAPDH R | GGCATGGACTGTGGTCATGAG |
| 18S F | GTAACCCGTTGAACCCCATT |
| 18S R | CCATCCAATCGGTAGTAGCG |
| RNase L F | GACACCTCTGCATAACGCAGT |
| RNase L R | AGGGCTTTGACCTTACCATACA |
| IFN-β F | GCCGCATTGACCATCTATGA |
| IFN-β R | GCCAGGAGGTTCTCAACAATAG |
| IL-6 F | ATCTAGATGCAATAACCACCCCT |
| IL-6 R | AGCTGCGCAGAATGAGATGA |
